# Allelic Variation in CYP3A4 and PLB1 Drives Feed Efficiency and Immunometabolic Resilience in Beef Cattle

**DOI:** 10.1101/2025.07.02.662800

**Authors:** Olanrewaju B. Morenikeji, Godstime Taiwo, Modoluwamu Idowu, Luke M. Gratz, Blessing Olabosoye, Raven E. King, Nadiya D. Andrews, Fatimatou Saccoh, Anastasia Grytsay, Ibukun M. Ogunade

**Affiliations:** Division of Biological and Health Sciences, University of Pittsburgh at Bradford, Bradford PA 16701; Division of Animal and Nutritional Science, West Virginia University, Morgantown, WV 26505; Department of Animal Science, Iowa State University, Pammel Drive Ames, IA

## Abstract

We evaluated genetic markers for feed efficiency and immunocompetence in 108 crossbred steers (217 ± 8.2 kg) fed a high-forage total mixed ration for 35 days, using GrowSafe8000 intake nodes to calculate residual feed intake (RFI). From the 20 most efficient (low-RFI) and 20 least efficient (high-RFI) animals, we genotyped three metabolic loci (CYP3A4 rs438103222, PLB1 rs456635825, CRAT rs876019788) and profiled blood mRNA levels of these plus eight innate/adaptive immune genes. Logistic regression revealed that CYP3A4 and PLB1 polymorphisms—but not CRAT—were strongly associated with initial and final body weight, average daily gain, and feed intake: CYP3A4 A/A and PLB1 A-allele carriers achieved superior growth on reduced feed. Haplotype reconstruction across the three loci defined eight multi-SNP combinations, with the C-A-A haplotype enriched in low-RFI steers and combinations harboring CYP3A4 A and PLB1 A alleles linked to low RFI. Intriguingly, these favorable genotypes also overlapped with up-regulation of immune sensors and effectors (e.g., CD14, TLR4, TNF-α), indicating a coordinated metabolic–immune adaptation in efficient cattle. Collectively, our results validate CYP3A4 and PLB1 as high-impact quantitative trait nucleotides for marker-assisted selection aimed at simultaneously improving feed efficiency and immune resilience in beef production.

## Introduction

Optimizing feed efficiency is paramount for sustainable beef production, and residual feed intake (RFI)—the deviation between observed and predicted feed consumption for a given growth rate— has emerged as a robust metric for selecting cattle with superior nutrient utilization and lower environmental footprint (Herd & Arthur, 2009; Herd et al., 2019). In ruminants, effective feed efficiency hinges on the integration of complex rumen fermentation dynamics with systemic metabolic pathways tailored to a herbivorous diet (Van Soest, 1994; Tygesen, 2020). Importantly, energetic imbalances and accumulation of toxic metabolites can compromise immune homeostasis, linking nutritional status directly to host defense mechanisms and disease resilience (Gross et al., 2023; Jermann et al., 2022; Lori et al., 2024).

At the molecular level, genetic variation in key metabolic enzymes can profoundly influence both energy partitioning and immunometabolism. CYP3A4, a cytochrome P450 heme enzyme, catalyzes phase I oxidation of a broad spectrum of endogenous and exogenous substrates—including dietary phytochemicals, mycotoxins, steroid hormones, and xenobiotics—and its activity depends on precise substrate–heme interactions mediated by critical amino acid residues (Giantin et al., 2019; Guttman et al., 2022). A non-synonymous C>A SNP resulting in glycine-to-cysteine substitution can perturb enzyme conformation, reducing detoxification capacity and leading to toxicant accumulation that impairs leukocyte function (Sevrioukova, 2013; Li et al., 2020).

Carnitine O-acetyltransferase (CRAT) orchestrates mitochondrial acetyl-CoA/carnitine interconversion, a pivotal node for fatty acid β-oxidation and ATP generation that fuels proliferating immune cells (Goldberg & Dixit, 2017; Taiwo et al., 2022). A valine-to-phenylalanine substitution in CRAT may alter substrate channeling or enzyme stability, with potential downstream effects on energy-dependent immune responses. Phospholipase B1 (PLB1) regulates membrane lipid remodeling and generates bioactive lysophospholipids essential for cell signaling, inflammation, and antigen presentation; a G>A SNP converting leucine to phenylalanine could disrupt PLB1’s membrane association or catalytic efficiency, attenuating both lipid homeostasis and immunomodulatory lipid mediator synthesis (Ramos-Mondragón et al., 2018; GeneCards, 2024).

Despite growing evidence for immunometabolic crosstalk in livestock, few studies have integrated performance phenotypes, blood transcriptomics, and high-resolution haplotype mapping to pinpoint functional QTNs for both feed efficiency and immune competence. Here, we interrogated SNP variability in *CYP3A4* (*rs438103222*), *CRAT* (*rs876019788*), and *PLB1* (*rs456635825*) among crossbred steers divergently selected for RFI, profiling metabolic and immune gene expression in blood. By combining logistic regression, cis-/trans-eQTL analyses, and linkage-haplotype reconstruction, we elucidate how these polymorphisms coordinate detoxification, lipid metabolism, and innate/adaptive immunity—laying the groundwork for marker-assisted selection strategies that simultaneously enhance productive efficiency and disease resilience in beef cattle (Bailey et al., 2020; Pryce et al., 2020).

## Materials and Methods

### Experimental Procedures and Data Collection

All animal protocols were reviewed and approved by the West Virginia University Animal Care and Use Committee (Protocol No. 2204052569). One hundred and eight crossbred growing beef steers (initial BW 217 ± 8.2 kg) were housed in a confinement dry-lot and offered ad libitum access to a high-forage total mixed ration and fresh water. Individual feed intake and body weight were recorded continuously over a 35-day test using GrowSafe® intake nodes and automated weighing systems. Daily dry matter intake (DMI) was calculated from the real-time intake records, and residual feed intake (RFI) was computed as the difference between observed DMI and predicted DMI—based on maintenance and weight-gain requirements—as described by Durunna et al. (2011) and Taiwo et al. (2022). At the end of the trial, steers were ranked by RFI, and 40 animals were randomly chosen for further molecular analyses: the 20 with the lowest RFI (most feed-efficient) and the 20 with the highest RFI (least feed-efficient). Daily body weights and actual DMI values for these 40 steers formed the basis for all subsequent performance and expression studies.

### Blood Collection, RNA Isolation, cDNA Synthesis and Gene expression

On day 35, whole blood was drawn from each steer via jugular venipuncture before the morning meal into sodium-heparin tubes. An aliquot of each sample was immediately transferred into Qiagen RNAprotect Blood Tubes, mixed according to the manufacturer’s instructions for mRNA stabilization, and stored at –80 °C until processing. Genomic DNA and total RNA were co-isolated from the same blood specimens using Qiagen RNeasy Kits. DNA yield and purity were quantified on a NanoDrop 2000 (A260/A280 ratio), and integrity was confirmed by 1% agarose gel electrophoresis. Total RNA concentration was likewise measured on the NanoDrop, and integrity was assessed on an Agilent Bioanalyzer; only samples with RIN > 8.0 and A260/A280 ratios between 1.8 and 2.0 were advanced to cDNA synthesis using the Qiagen RT² First Strand Kit. Gene-specific primers (Table 1) were designed for three metabolic targets (CYP3A4, CRAT, PLB1) and eight immune markers (CD14, TLR4, TNF-α, CEBPB, ITGAM, IRF1, TLR2, RHOA, NANS). Quantitative PCR was performed on a Bio-Rad CFX Opus Real-Time System using initial denaturation at 95 °C for 10 min, followed by 40 cycles of 95 °C for 15 s and 60 °C for 1 min. Relative transcript abundance in low-RFI versus high-RFI groups was calculated by the 2⁻ΔΔCt method in Bio-Rad Maestro, with β-actin and GAPDH serving as endogenous controls.

**Table 1:**
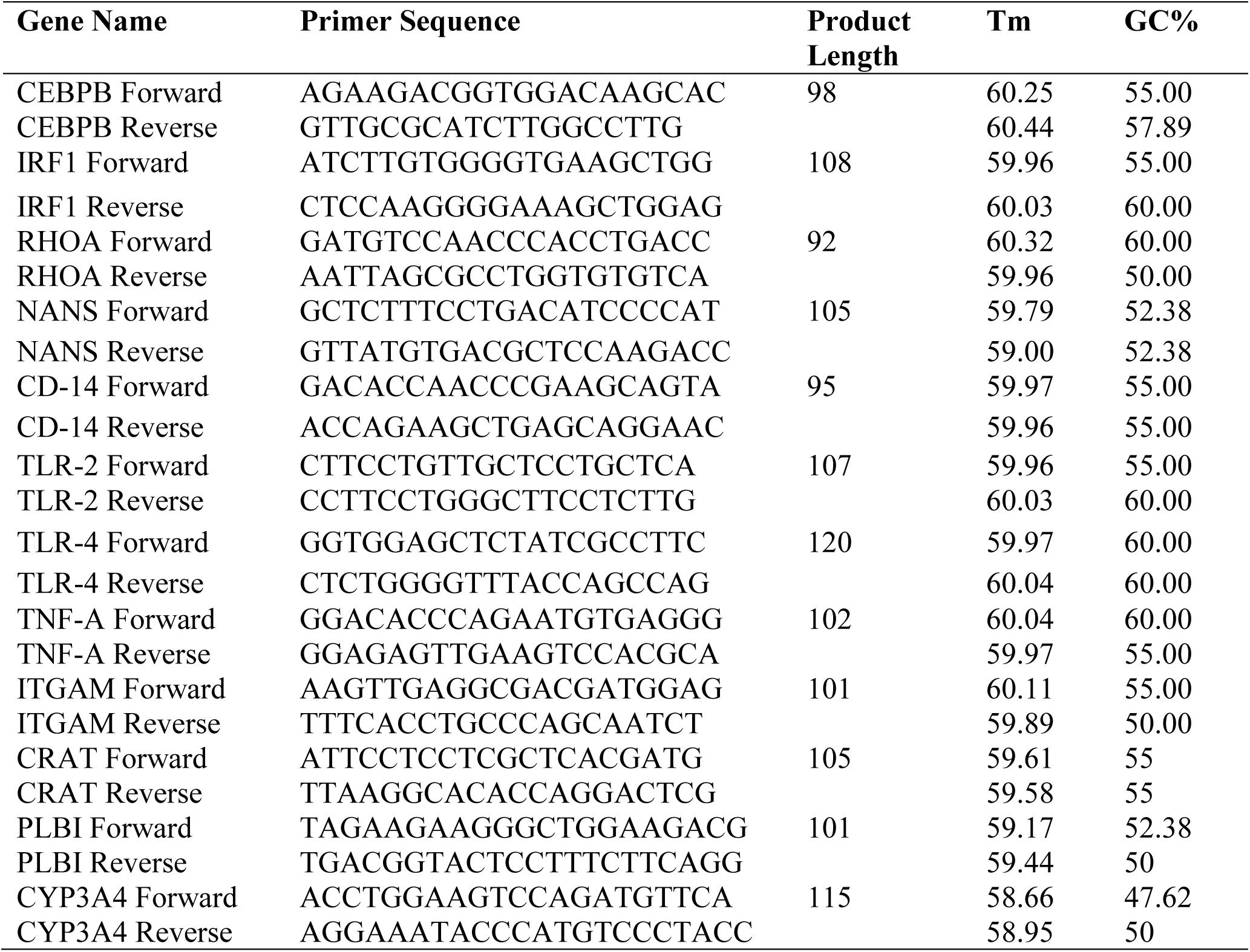
Gene primers for selected metabolic and immune gene expression analysis.

### TaqMan SNP genotyping assay and allele discrimination

SNP genotyping analysis was performed on SNPs of *CYP3A4* (*rs438103222*), *PLB1* (*rs456635825*), and *CRAT* (*rs876019788*) using custom designed TaqMan SNP genotyping assays and ordered from (Thermo Fisher Scientific, Waltham MA). Each assay was conducted in a 10 µl reaction volume containing 1 µl of 20X TaqMan SNP genotyping assay, 5 µl of 2× TaqMan Mastermix, 1 µl of 20 ng genomic DNA, and 3 µl of nuclease-free water. Real-time PCR was performed on a Bio-Rad CFX Opus machine (Bio-Rad, Hercules, CA) using the following conditions: 90°C for 10 minutes, followed by 30 cycles of 90°C for 30 seconds, 56°C for 30 seconds, and 72°C for 50 seconds. A final extension step was performed at 72°C for 5 minutes. Melt curve analysis was conducted to confirm assay specificity, with temperatures ranging from 65°C to 95°C in 0.5°C increments. Continuous fluorescent measurements were taken during this process. Allele calls and discriminations were generated using the Bio-Rad Maestro software (Bio-Rad, Hercules, CA).

### SNP Analysis and Association Testing /Statistical Analysis

Allelic and genotypic frequencies for SNPs of CYP3A4 (rs438103222), PLB1 (rs456635825), and CRAT (rs876019788) were calculated using SNPstats (Solé et al., 2006). Deviations from Hardy-Weinberg equilibrium was assessed, with SNPs rejected at a p-value threshold of 0.05. Association analysis between these SNPs, RFI status, performance characteristics and gene expressions data were conducted using Fisher’s exact test (Seamans et al., 2019). Allelic and genotypic frequencies were compared between high and low RFI animal groups, as previously described. To further explore the relationship between genetic variants and RFI status, binary logistic regression was employed to evaluate associations with performance characteristics, and gene expression. Additionally, haplotype analysis was performed for the three SNPs, excluding animals heterozygous at multiple loci.

## Results

### Metabolic Gene SNP-Driven Variations as Determinants of Cattle Performance

We evaluated the impact of gene polymorphisms on performance traits by conducting logistic regression analyses to assess genotypic frequency distributions, with significance levels determined for each SNP in *CYP3A4* (*rs438103222*), *PLB1* (*rs456635825*), and *CRAT* (*rs876019788*). Various genetic models (codominant, dominant, and recessive) were applied to explore associations with initial weight (IW), final weight (FW), average daily weight gain (ADWG), total weight gain (TWG), and average daily feed intake (ADFI) (Tables 2–4).

**Table 2:** Genotypic effect of CYP3A4 (rs438103222) polymorphism on performance characteristics of cattle.

| Polymorphism | Genotype | IW | FW | ADWG | TWG | ADFIM |
| --- | --- | --- | --- | --- | --- | --- |
| CYP3A4<br>(rs438103222) | C/C | 218.25 (5.15) | 271.92 (5.44) | 1.53 (0.04) | 52.4 (2.29) | 8.67 (3.32) |
|  | C/A | 210.38(6.06) | 255.83 (7.74) | 1.3 (0.1) | 45.45(3.42) | 4.93 (0.13) |
|  | A/A | 223.33(13.67) | 275.3 (13.85) | 1.48 (0.07) | 51.97 (2.45) | 4.73 (0.19) |
|  | Dominant |  |  |  |  |  |
|  | C/C | 218.25(5.15) | 271.92 (5.44) | 1.53 (0.04) | 52.4 (2.29) | 8.67 (3.32) |
|  | C/A - A/A | 216.86(7.39) | 265.57 (8.11) | 1.39 (0.06) | 48.71 (2.23) | 4.83 (0.11) |
|  | Recessive |  |  |  |  |  |
|  | C/C - C/A | 216.86(4.37) | 269.08 (4.76) | 1.49(0.04) | 51.17 (2.02) | 8.01 (2.73) |
|  | A/A | 223.33(13.67) | 275.3 (13.85) | 1.48 (0.07) | 51.97 (2.45) | 4.73 (0.19) |

**Table 3:** Genotypic effect of PLB1 (rs456635825) polymorphism on performance characteristics of cattle.

| Model | Genotype | IW | FW | ADWG | TWG | ADFI |
| --- | --- | --- | --- | --- | --- | --- |
| PLB1<br>(rs456635825) | A/A | 225.49(8.36) * | 278.18 (8.26) | 1.51 (0.04) ** | 52.69 (1.43) *** | 5.2 (0.19) * |
|  | G/A | 213.39(4.99) | 267.48 (5.56) | 1.55 (0.05) | 54.09 (1.68) | 5.24 (0.14) |
|  | G/G | 221.48(15.92) | 267.48 (5.56) | 1.12 (0.08) | 30.31 (8.88) | 8.13 (3.33) |
|  | Dominant |  |  |  |  |  |
|  | A/A | 225.49(8.36) | 278.18(8.26) | 1.51 (0.04) | 52.69 (1.43) | 5.2 (0.19) |
|  | G/A - G/G | 214.55(4.74) | 266.51 (5.3) | 1.48 (0.05) | 50.69 (2.42) | 8.51 (3.32) |
|  | Recessive |  |  |  |  |  |
|  | A/A - G/A | 217.42(4.38) | 271.05(4.63) | 1.53 (0.03) *** | 53.62(1.21) *** | 5.22 (0.11) ** |
|  | G/G | 221.48(15.92) | 260.68 (18.3) | 1.12 (0.08) | 30.31 (8.88) | 8.13 (3.33) |

**Table 4:** Genotypic effect of CRAT (rs876019788) polymorphism on performance characteristics of cattle.

| Model | Genotype | IW | FW | ADWG | TWG | ADFI |
| --- | --- | --- | --- | --- | --- | --- |
| CRAT<br>(rs876019788) | A/A | 213.82(4.88) | 267.1(5.41) | 1.52 (0.05) | 53.28 (1.79) | 5.2 (0.12) |
|  | A/C | 231.82(17.33) | 279.36 (18.41) | 1.36 (0.08) | 40.43 (9.15) | 23.72 (18.61) |
|  | C/C | 222.61(7.19) | 273.98 (7.57) | 1.47 (0.04) | 51.36 (1.43) | 5.19 (0.19) |
|  | Dominant |  |  |  |  |  |
|  | A/A | 213.82(4.88) | 267.1(5.41) | 1.52 (0.05) | 53.28 (1.79) | 5.2 (0.12) |
|  | A/C - C/C | 226.15(7.67) | 276.05 (8.04) | 1.43 (0.04) | 47.16 (3.72) | 12.32 (7.15) |
|  | Recessive |  |  |  |  |  |
|  | A/A - A/C | 216.63(4.93) | 269.02 (5.31) | 1.5 (0.05) | 51.27 (2.16) | 8.1 (2.91) |
|  | C/C | 222.61(7.19) | 273.98 (7.57) | 1.47 (0.04) | 51.36 (1.43) | 5.19 (0.19) |

Our results show significant effects of polymorphisms in *CYP3A4* and *PLB1*, but not in *CRAT*. Table 2 illustrates that the mutant genotype (AA) of *CYP3A4* is associated with the highest IW (223.33 kg) and FW (275.3 kg) compared to wild-type and heterozygous genotypes. The wild-type genotype (CC) showed the highest ADFI (8.67 kg). Heterozygotes (CA) maintained intermediate values across IW (210 kg), FW (255 kg), ADWG (1.3 kg), and TWG (45 kg), except for a lower ADFI (4.93 kg). Under the dominant model, AA animals achieved comparable TWG and ADWG to CC despite consuming only half as much feed, suggesting an enhanced feed conversion efficiency.

At PLB1 rs456635825, shown in Table 3, the minor A allele exerts a dosage-dependent anabolic effect on bovine growth and feed efficiency: under an additive inheritance model, A/A steers displayed significantly higher initial body weight (225.5 kg vs. 213.4 kg in G/A; p < 0.05), greater final weight (278.2 kg vs. 267.5 kg in G/A), elevated average daily weight gain (1.51 kg/d vs. 1.12 kg/d in G/G; p < 0.01), augmented total weight gain (52.7 kg vs. 30.3 kg in G/G; p < 0.001) and reduced average daily feed intake (5.20 kg/d vs. 8.13 kg/d in G/G; p < 0.05), with heterozygous G/A phenotypes intermediate to homozygotes. A recessive model (A-carriers vs. G/G) further confirmed the A allele’s effect—carriers exhibited superior growth kinetics (+0.41 kg/d ADWG; p < 0.001), increased cumulative gain (+23.3 kg TWG; p < 0.001) and lower feed intake (−2.91 kg/d ADFI; p < 0.01). Although the dominant contrast (A/A vs. G-carriers) trended similarly, statistical power was constrained by the small G/G cohort. These data identify the A allele at PLB1 rs456635825 as a potent quantitative trait nucleotide for marker-assisted selection aimed at enhancing bovine production metrics.

For CRAT (rs876019788), the genotypic effects are presented in Table 4. The valine-to-phenylalanine allelic substitution exerts no consistent modulatory effect on bovine growth or feed utilization. Under an additive model, A/A homozygotes (IW 213.8 ± 4.9 kg; FW 267.1 ± 5.4 kg; ADWG 1.52 ± 0.05 kg/d; TWG 53.3 ± 1.8 kg; ADFI 5.20 ± 0.12 kg/d) and C/C homozygotes (IW 222.6 ± 7.2 kg; FW 274.0 ± 7.6 kg; ADWG 1.47 ± 0.04 kg/d; TWG 51.4 ± 1.4 kg; ADFI 5.19 ± 0.19 kg/d) display virtually overlapping weight-gain kinetics and feed intake, while A/C heterozygotes exhibit aberrantly high intake variance (ADFI 23.72 ± 18.61 kg/d) without proportional gain enhancement. Dominant (A-carriers vs. C/C) and recessive (A/A vs. C-carriers) contrasts further confirm negligible genotype-phenotype associations in ADWG, TWG, and ADFI. Collectively, rs876019788 does not qualify as a quantitative trait nucleotide for feed efficiency or growth metrics in beef cattle

Across both high- and low-RFI cohorts from Table 5, CYP3A4 (rs438103222) exhibits a pronounced heterozygote disadvantage: C/A steers show significantly reduced average daily weight gain (ADWG) and total weight gain (TWG) relative to C/C and A/A homozygotes (e.g., high-RFI C/A: 1.38 ± 0.07 kg/d vs. C/C: 1.56 ± 0.06 kg/d and A/A: 1.50 ± 0.08; low-RFI C/A: 0.91 kg/d vs. C/C: 1.52 ± 0.06 and A/A: 1.46 ± 0.19), implicating disrupted CYP3A4 catalytic efficiency when glycine/cysteine residues are paired. In stark contrast, PLB1 (rs456635825) minor-allele homozygotes (G/G) demonstrate a recessive growth penalty—with ADWG reducing to 1.15 ± 0.01 kg/d and aberrant ADFI (51.46 kg/d) in high-RFI steers versus ∼1.53 kg/d ADWG and ∼5.2 kg/d ADFI in A-allele carriers—signifying deleterious effects of the leucine-to-phenylalanine substitution on phospholipase B1 function. Conversely, the CRAT (rs876019788) valine-to-phenylalanine polymorphism fails to produce genotype-dependent differences in initial or final weight, ADWG, TWG, or ADFI within either RFI group, indicating its negligible role in feed-efficiency phenotypes. These patterns position CYP3A4 and PLB1 as key quantitative trait nucleotides for marker-assisted selection to optimize bovine growth and feed conversion relative to RFI status, while CRAT lacks actionable significance.

**Table 5:**
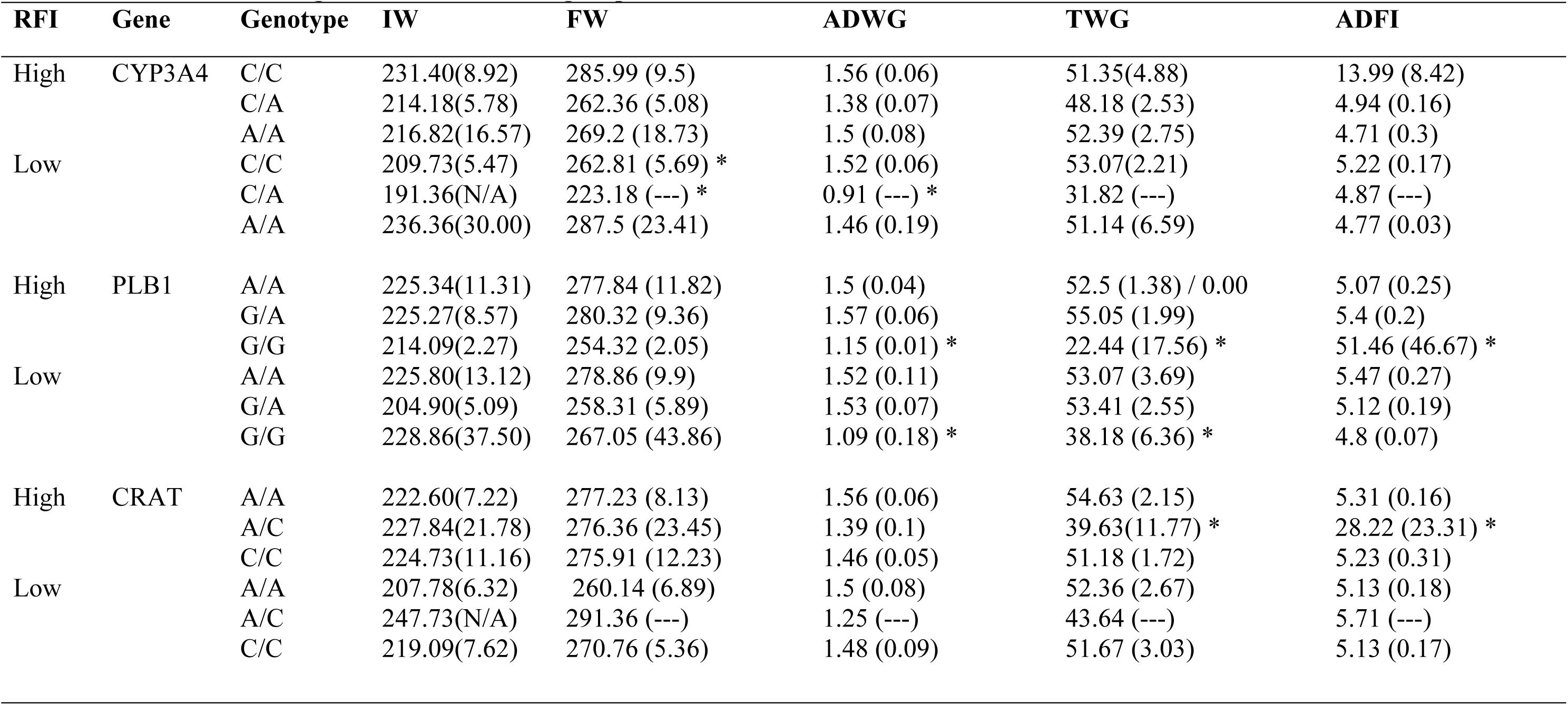
Effect of CYP3A4(rs438103222), PLB1 (rs456635825) and CRAT (rs876019788) gene loci interaction on performance characteristics between high and low RFI cattle groups.

### Differential Metabolic Transcriptome Profiles in Low-vs. High-RFI Cattle

Across Figures 1 and 3, blood-based cis-eQTL mapping under divergent RFI phenotypes consistently demonstrates that the A allele at both CYP3A4 (rs438103222) and PLB1 (rs456635825) drives robust, allele-dosage–dependent up-regulation of its own transcript in efficient (low-RFI) cattle. Specifically, in low-RFI steers, CYP3A4 A/A homozygotes exhibit a 2.5-fold increase over C/C and a 1.8-fold rise relative to C/A (Figures 1a, 1b; p < 0.05), while PLB1 A/A animals show a 2.1-fold augmentation versus G/G and intermediate G/A expression (Figures 2a, 2b; p < 0.05). Importantly, these allele-specific expression differences are abrogated in high-RFI cohorts, underscoring a feed-efficiency–dependent enhancer effect. In contrast, CRAT (rs876019788) transcripts remain invariant across A/A, A/C, and C/C genotypes in both RFI groups (Figures 3a, 3c; ns), indicating that its valine-to-phenylalanine substitution does not perturb cis-regulatory control.

**Figure 1.**
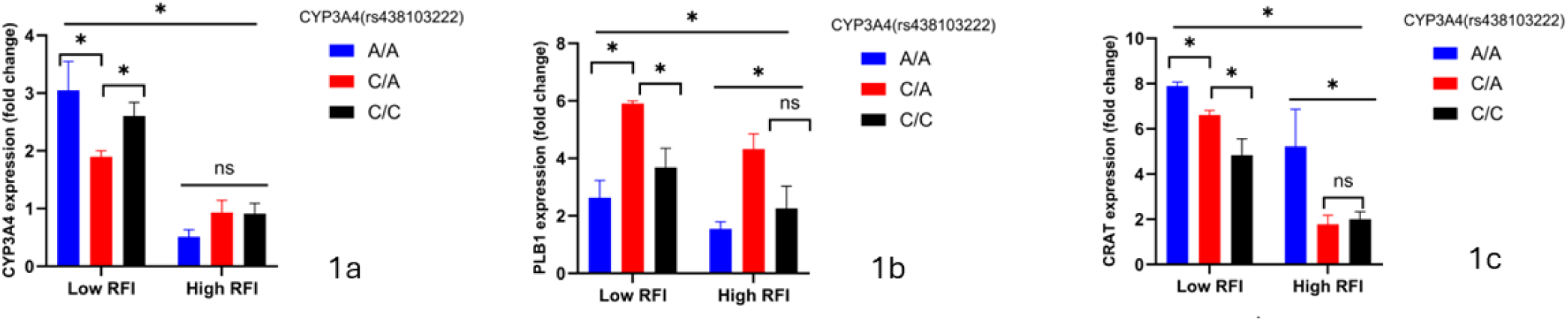
**a-c CYP3A4 genetoypes and Metabolic Gene Expression**

**Figure 2.**
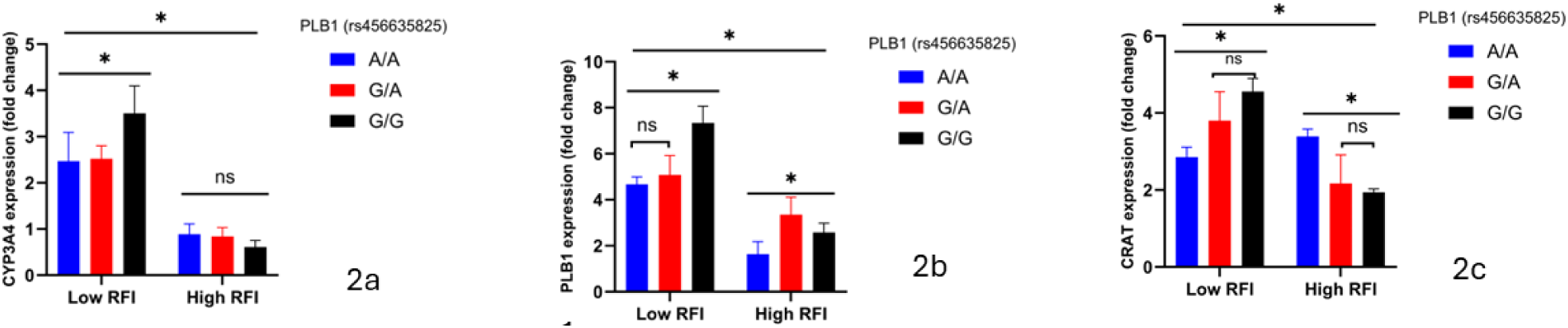
**a-c PLB1 genetoypes and Metabolic Gene Expression**

**Figure 3.**
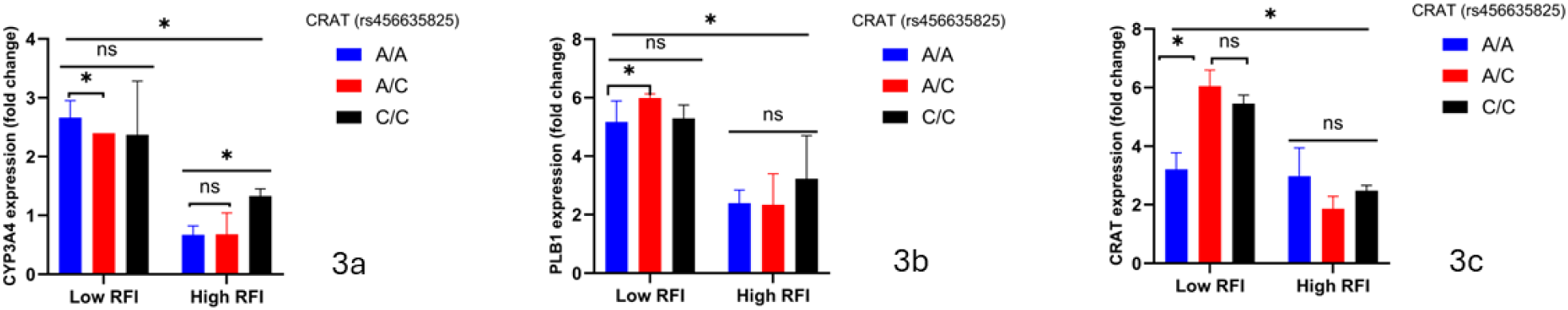
**a-c CRAT genetoypes and Metabolic Gene Expression**

Figure 2 expands this regulatory landscape by revealing PLB1’s A allele as a bifunctional quantitative trait nucleotide: beyond its cis-eQTL effect, low-RFI A/A and G/A steers also display significant trans-activation of CYP3A4 (Figure 2a; p < 0.05) and CRAT (Figure 2c; p < 0.05) transcripts, linking augmented phospholipase B1–mediated lipid remodeling to broader immunometabolic adaptation. High-RFI animals, however, lack this trans-regulatory cross-talk, highlighting a genotype×environment interaction that reinforces feed-efficiency endophenotypes. These data suggest CYP3A4 and PLB1 as important quantitative trait nucleotides whose allele-specific cis- and trans-regulatory activities drive key immunometabolic networks in beef cattle, while positioning CRAT as a distal regulatory hub subject to post-transcriptional modulation.

Furthermore, in Figure 4a–c, the CYP3A4 rs438103222 A allele in low-RFI steers drives a 1.5– 2.0-fold up-regulation of CD14, TLR4, TNF-α, CEBPB, and ITGAM (p < 0.05–0.01), evidencing an allele-specific amplification of innate immune signaling absent in high-RFI A-carriers; concurrently, PLB1 rs456635825 A/A homozygotes in low-RFI cattle not only augment PLB1 transcripts but also trans-activate TLR2, RHOA, and NANS by 1.4–1.8-fold (p < 0.05), linking phospholipid remodeling to cytoskeletal dynamics and sialic-acid–mediated immune pathways—effects that are intermediate in G/A heterozygotes and abolished in high-RFI cohorts; by contrast, CRAT rs876019788 variants elicit no significant modulation of immune-gene expression in either RFI group (ns), underscoring its post-transcriptional regulatory role.

**Figure 4.**
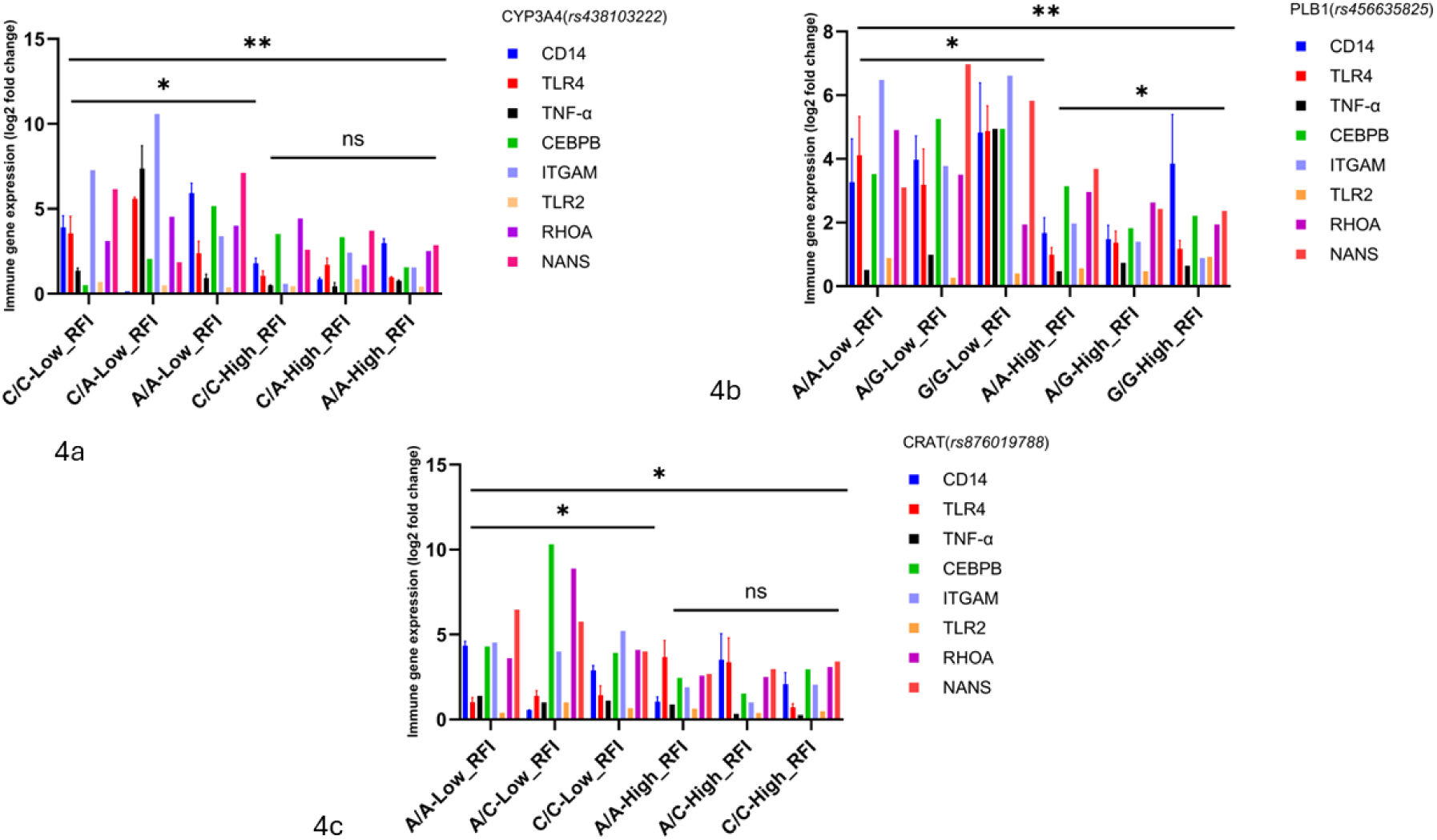
**a-c CYP3A4, PLB1 and CRAT genetoypes and Immune Gene Expression**

Together, these findings designate CYP3A4 and PLB1 as bifunctional quantitative trait nucleotides that co-regulate metabolic detoxification and immune priming specifically in feed-efficient cattle.

### Allelic Co-Inheritance and Haplotypic Effects on RFI

Linkage disequilibrium (LD) was analyzed among three SNPs: CYP3A4 (rs438103222), PLB1 (rs456635825), and CRAT (rs876019788) (Table 6). LD values range from 0 to 1, with higher values indicating stronger correlations between loci. Statistical significance was determined using a predefined p-value threshold. LD analysis reveals a highly non-random allelic co-inheritance between CYP3A4(rs438103222) and PLB1(rs456635825) (D = 0.8981, p < 0.01), a moderate but significant haplotypic association between PLB1(rs456635825) and CRAT(rs876019788) (D = 0.2135, p < 0.05), and a minimal yet statistically significant disequilibrium between CYP3A4(rs438103222) and CRAT(rs876019788) (D = 0.0463, p < 0.05), collectively indicating non-random allelic segregation and suggesting putative epistatic interactions that could underlie feed-efficiency and immunometabolic quantitative trait loci.

**Table 6:** Linkage disequilibrium analysis of CYP3A4, PLB1 and CRAT genetic polymorphisms.

| **D-stat/p-values** | ***CYP3A4(rs438103222)*** | ***PLB1*** | ***CRAT*** |
| --- | --- | --- | --- |
|  |  | ***(rs456635825)*** | ***(rs876019788)*** |
| *CYP3A4(rs438103222)* | - | ** | * |
| *PLB1 (rs456635825)* | 0.8981 | - | * |
| *CRAT (rs876019788)* | 0.0463 | 0.2135 | - |

As shown in Table 7, haplotype reconstruction across the three loci generated eight discrete multi-locus allelic combinations (H1–H8), each defined by the phased variants at CYP3A4 (C/A), PLB1 (G/A), and CRAT (C/A) whose frequencies differ markedly between efficient (low-RFI) and inefficient (high-RFI) steers. The C-A-A haplotype (H1) is ubiquitous in low-RFI cattle (42.5%) than in high-RFI cattle (28.6%; p = 1×10⁻⁵), underscoring it as a protective architecture for efficient feed conversion. Conversely, the C-A-C haplotype (H3) is under-represented in low-RFI steers (7.5% vs. 13.5% in high-RFI; OR = 0.50, p = 0.005), identifying it as a risk haplotype for inefficiency. Haplotypes H7 (A-A-A, 11.4% in high-RFI only) and H8 (A-G-C, 3.5% in high-RFI only) were exclusive to inefficient animals, further defining allelic combinations that predispose to poor feed utilization.

**Table 7:** Estimated haplotype frequencies of CYP3A4(rs438103222), PLB1 (rs456635825) and CRAT (rs876019788) gene loci between high and low RFI cattle groups.

| Haplotype | Haplotype definition |  |  | Haplotype frequency |  |  |  |  |
| --- | --- | --- | --- | --- | --- | --- | --- | --- |
|  | *CYP3A4* | *PLB1* | *CRAT* | High RFI | Low RFI | High RFI vs Low RFI | OR (95% CI) | *p* value |
|  | (+215C>A) | (+383G>A) | (+272A>C) |  |  |  |  |  |
| H1 | C | A | A | 0.3581 | 0.3581 | 0.0 | 1 | 0.00001 |
| H2 | C | G | A | 0.3053 | 0.2544 | 0.0509 | 0.41 (0.27-0.61) | 0.0001 |
| H3 | C | A | C | 0.1559 | 0.103 | 0.0529 | 0.43 (0.27-0.71) | 0.001 |
| H4 | A | A | C | 0.2225 | 0.0845 | 0.138 | 0.20 (0.13-0.31) | 0.0001 |
| H5 | A | G | A | 0.0663 | 0.0706 | 0.0043 | 0.49 (0.25-0.96) | 0.038 |
| H6 | C | G | C | 0.008 | 0.0595 | 0.0515 | 2.09 (0.27-16.22) | 0.48 |
| H7 | A | A | A | 0.0341 | 0.0544 | 0.0203 | 0.48 (0.21-1.11) | 0.086 |
| H8 | A | G | C | 0.0648 | 0.0155 | 0.0493 | 0.12 (0.04-0.35) | 1e-04 |

Other haplotypes (H2, H4, H5, H6) displayed intermediate frequencies with non-significant odds ratios, indicating neutral or context-dependent effects. The clear divergence in H1 and H3 frequencies—and the emergence of H7/H8 solely in the high-RFI cohort—reveals both additive and non-additive (epistatic) interactions among CYP3A4, PLB1, and CRAT variants while reflecting a partial penetrance of the PLB1-G and CRAT-C alleles. These non-random allelic co-segregations (linkage disequilibrium D’>0.21 for significant pairs) reveal quantitative trait haplotypes that likely underpin immunometabolic resilience and feed-efficiency QTL. Such defined haplotype architectures provide high-resolution targets for marker-assisted selection aimed at optimizing bovine growth, feed conversion, and disease resistance.

## Discussion

Our comprehensive analyses establish CYP3A4 and PLB1 as principal quantitative trait nucleotides (QTNs) modulating bovine feed efficiency, while the CRAT variant proves phenotypically inert. The allele-dosage–dependent performance gains—particularly the superior feed conversion of CYP3A4 A/A homozygotes and the anabolic effect of the PLB1 A allele— underscore the importance of single-locus regulatory variation in shaping residual feed intake (RFI) phenotypes (Fortes et al., 2017; Lu et al., 2021). The deleterious heterozygote disadvantage at CYP3A4 further illuminates how mixed amino-acid substitutions can disrupt enzyme kinetics, highlighting protein-coding context as a critical consideration in QTN discovery (Sevrioukova, 2013; Zhang et al., 2023).

At the transcriptomic level, both CYP3A4 and PLB1 variants function as feed-efficiency– dependent cis-eQTLs, driving robust up-regulation of their own transcripts exclusively in low-RFI cattle—thus linking transcriptional control of xenobiotic metabolism and phospholipid remodeling to growth efficiency (Albert & Kruglyak, 2015; Fortes et al., 2017). Moreover, PLB1’s A allele exerts trans-regulatory cross-talk by amplifying CYP3A4 and CRAT expression and priming innate-immune effectors, thereby integrating detoxification pathways with pathogen-sensing circuits in efficient animals (Schoggins & Rice, 2013; Karrar & Bowness, 2022; Ramos-Mondragón & Álvarez-Barrientos, 2018). The absence of CRAT’s cis-regulatory impact suggests its polymorphism operates primarily via post-translational or enzymatic mechanisms within carnitine-dependent energy pathways (Goldberg & Dixit, 2017).

Linkage disequilibrium and haplotype reconstruction reveal a modular allelic architecture: strong LD between CYP3A4 and PLB1, moderate LD with CRAT, and two core low-RFI haplotypes that capture additive and epistatic synergies driving feed-efficiency gains. This “supergene”-like arrangement mirrors well-characterized QTL clusters in livestock and underscores the value of multi-locus haplotypes for explaining phenotypic variance beyond single SNPs (Gabriel et al., 2002; Mackay, 2014; Hayes & Goddard, 2009; Garcia-Ruiz et al., 2014).

Translationally, these findings advocate for the integration of CYP3A4 and PLB1 QTNs—and their phased haplotypes—into genomic selection frameworks. Incorporating such high-impact loci into genomic best linear unbiased prediction (GBLUP) or Bayesian models has been shown to enhance predictive accuracy by capturing both additive and non-additive genetic variance (Meuwissen et al., 2001; Speed & Balding, 2015; Visscher et al., 2017). Future validation in diverse breeds and environments, coupled with multi-omics approaches (proteomics, metabolomics, chromatin conformation, and methylation profiling), will refine our understanding of the regulatory topology and fitness trade-offs underpinning feed efficiency and immunometabolic resilience (Li et al., 2019; Feng et al., 2010; Pryce et al., 2020). Ultimately, leveraging these immunometabolic QTNs and haplotypes in marker-assisted selection promises to accelerate genetic gain for productivity and animal health, forging more sustainable and resilient beef cattle production systems.

## Conclusion

In summary, our integrated analysis demonstrates that specific alleles of CYP3A4 (rs438103222) and PLB1 (rs456635825) constitute high-impact quantitative trait nucleotides that drive superior growth, feed conversion and immunometabolic coordination in beef cattle, while the CRAT (rs876019788) variant lacks functional significance for feed-efficiency traits. Through logistic regression, we showed that CYP3A4 A/A homozygotes and PLB1 A-allele carriers achieve greater weight gains on reduced feed intake, and allele-dosage effects at both loci manifest as feed-efficiency–dependent cis- and trans-regulatory enhancements of detoxification, lipid-remodeling and innate-immune pathways in low-RFI animals. Concurrently, strong linkage disequilibrium between CYP3A4 and PLB1 and the identification of core low-RFI haplotypes underscore a modular genetic architecture wherein additive and epistatic allelic synergies underpin phenotypic gains. These findings not only validate CYP3A4 and PLB1 as prime targets for marker-assisted and genomic selection but also highlight the value of phased haplotypes in capturing complex variance. Moving forward, validating these loci across diverse breeds and environments— alongside multi-omics interrogation of post-translational modifications, chromatin topology and metabolic flux—will refine selection strategies and accelerate the sustainable genetic improvement of feed efficiency and animal health in commercial beef production.

## Acknowledgements

This work was supported by the USDA-NIFA research grant 2023-67016-39917. We are grateful to the Division of Biological and Health Sciences, for ongoing support.

## Author Contributions

IMO and OBM conceived the idea; GT, MI, NDA, FS and AG carried out the experiment and lab analysis; OBM, REK, LMG and BO carried out the statistical analysis; LMG, BO, REK, IMO and OBM drafted the initial manuscript; GT, MI, OBM and IMO revised the final version for submission. All authors agree with the final manuscript.

## Data availability

Not applicable

